## Supplementary material for "Arms races between selfish genetic elements and their host defence": Qiu et al_Suppl Material_rev_final.docx

**Supplementary Table 1. Sample information and statistics for the genome assemblies**

| Assembly | *Mastotermes darwiniesis* | *Zootermopsis nevadensis* | *Cryptotermes secundus* | *Reticulitermes grassei* | *Trinervitermes geminatus ** | *Odontotermes sp.2* | *Macrotermes bellicosus (Q)* | *Macrotermes bellicosus (S)* |
| --- | --- | --- | --- | --- | --- | --- | --- | --- |
| Caste | Worker | Worker | Worker | Queen | Worker | Queen | Queen | Soldier |
| Sample age # | 1-2 years | 1-2 years | 3-4 years | > 3 years | Few months | 4 years | > 7 years | Few months |
| Library | HiFi standard | HiFi standard | HiFi low input | HiFi standard | HiFi low input | HiFi low input | HiFi standard | HiFi low input |
| Tissue | Head | Head | Gut-removed whole bodies | Gut-removed whole bodies | Gut-removed whole bodies | Fatbody | Fatbody | Gut-removed whole bodies |
| # contigs | 2,206 | 321 | 3,582 | 98 | 3,206 | 1234 | 364 | 6,173 |
| Assembly length (bp) | 1,345,564,086 | 576,263,831 | 1,304,526,754 | 996,825,901 | 1,453,058,254 | 1,465,002,929 | 1,340,725,563 | 1,428,941,907 |
| GC (%) | 39.53 | 38.48 | 41.18 | 39.49 | 40.36 | 39.84 | 40.26 | 40.31 |
| Largest contig (bp) | 14,453,904 | 27,517,609 | 7,090,375 | 72,273,710 | 6,339,778 | 22,652,318 | 50,731,358 | 2,705,196 |
| N50 (bp) | 1,569,491 | 18,733,084 | 1,070,193 | 44,444,803 | 809,495 | 6,388,291 | 15,380,714 | 528,038 |
| L50 | 212 | 13 | 353 | 10 | 520 | 73 | 28 | 813 |
| **PacBio data statistics** | | | | | | | | |
| Total reads | 1,870,375 | 2,151,234 | 2,483,158 | 2,328,724 | 1,994,195 | 4,418,500 | 1,773,689 | 3,613,653 |
| Raw base (Gb) | 17.4 | 29.9 | 17.7 | 27.6 | 19.2 | 39.8 | 29.6 | 18.2 |
| Mean read length (bp) | 9,303 | 13,474 | 7,139 | 11,836 | 9,598 | 9,010 | 16,689 | 5,028 |
| Read coverage | 22 | 66 | 18 | 30 | 16 | 33 | 27 | 16 |
| **GenomeScope results** (k-mer = 21) | | | | | | | | |
| Est. haploid genome length (bp) | 1,317,818,343 | 561,384,891 | 1,142,543,631 | 950,124,871 | 1,226,968,067 | 1,281,361,698 | 1,273,604,650 | 1,282,772,194 |
| Est. heterozygosity (%) | 0.22 | 0.45 | 5.27 | 0.23 | 1.2 | 0.79 | 0.35 | 0.39 |
| Est. repeat length (bp) | 547,115,452 | 173,947,255 | 591,445,906q | 289,378,874 | 858,067,778 | 526,497,403 | 499,855,941 | 502,589,585 |
| **BUSCO results** | | | | | | | | |
| Completeness | 1364 (99.8%) | 1367 (100.0%) | 1364 (99.8%) | 1364 (99.8%) | Pre:  1357 (99.3%)  Post: 1351 (98.8%) | 1363 (99.8%) | 1365 (99.9%) | 1363 (99.8%) |
| Single-copy | 1293 (94.6%) | 1353 (99.0%) | 1206 (88.2%) | 1348 (98.6%) | Pre:  1161 (84.9%)  Post: 1315 (96.2%) | 1339 (98.0%) | 1316 (96.3%) | 1231 (90.1%) |
| Duplicated | 71 (5.2%) | 14 (1%) | 158 (11.6%) | 16 (1.2%) | Pre:  196 (14.3%)  Post: 36 (2.6%) | 24 (1.8%) | 49 (3.6%) | 132 (9.7%) |
| Fragmented | 1 (0.1%) | 0 (0%) | 3 (0.2%) | 2 (0.1%) | Pre:  7 (0.5%)  Post: 7 (0.5%) | 3 (0.2%) | 1 (0.1%) | 3 (0.2%) |
| Missing | 2 (0.2%) | 0 (0%) | 0 (0%) | 1 (0.1%) | Pre:  3 (0.2%)  Post: 9 (0.7%) | 1 (0.1%) | 1 (0.1%) | 1 (0.1%) |
| **Merqury results** (k-mer = 21) | | | | | | | | |
| K-mer completeness | 98.6 % | 95.6 % | 91.2 % | 97.3 % | Pre: 96.1 %  Post: 93.9 % | 93.2 % | 97.2 % | 96.8 % |
| Assembly error rates | 8x10^-7^ | 2x10^-7^ | 3x10^-6^ | 1x10^-7^ | Pre: 5x10^-6^  Post: 3x10^-6^ | 1x10^-6^ | 9x10^-7^ | 2x10^-6^ |
| QV | 61 | 66 | 55 | 69 | Pre: 53  Post: 57 | 60 | 60 | 57 |

**^*^** For *Trinervitermes geminatus*, the genome assembly has been processed with *purge_dups* to remove potential haplotypic duplications. BUSCO and Merqury results are provided for both the pre- and post-purging assemblies.

^#^ Although there was considerable variation in chronological age among the individuals used, they were of intermediate age relative to their species and caste identity.

**Supplementary Table 2. Abundances of TE families (a) and superfamilies (b) and in termite and woodroach genomes.** As separate excel file.

**Supplementary Table 3. CpG methylation levels of TE superfamilies in termite genomes.** As separate excel file.

**Supplementary Table 4. TE defense genes in termite and woodroach genomes.** As separate excel file.

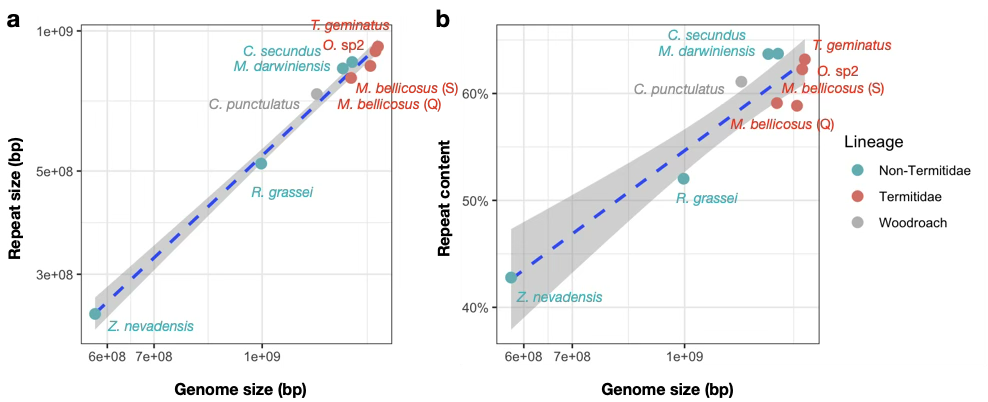

**Supplementary Figure 1.** Genome size of termites and the woodroach plotted against (**a**) genomic repeat size and (**b**) the proportion of repeat content (percentage of genome size). Both genomic repeat size and repeat content proportion show significant positive correlations with genome size (Pearson's correlation tests; both *p* < 1×10⁻³).

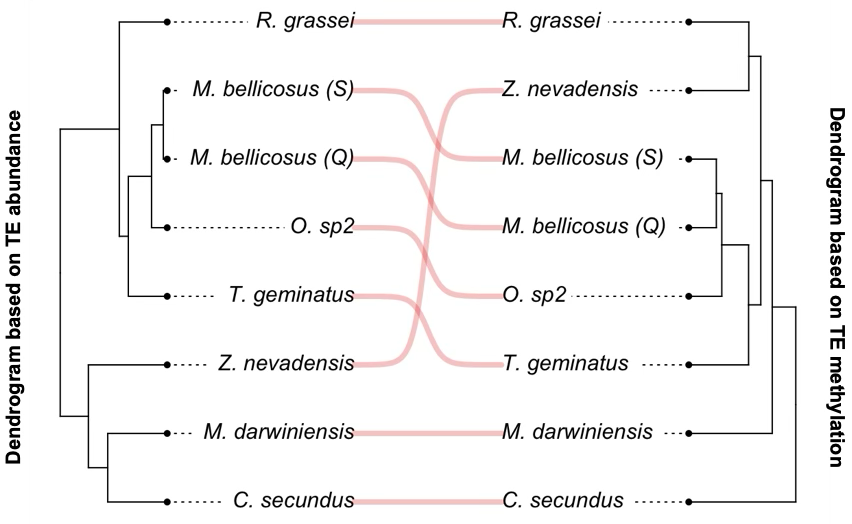

**Supplementary Figure 2.** Dendrogram based on TE superfamily abundance (*n_superfamily_* = 71) (left) and on TE superfamily methylation (*n_superfamily_* = 69) (right) across the termite phylogeny. TE abundance and TE methylation show a strong co-phylogenetic pattern (permutation test; *n _permutations_* = 100, *p* < 0.01).

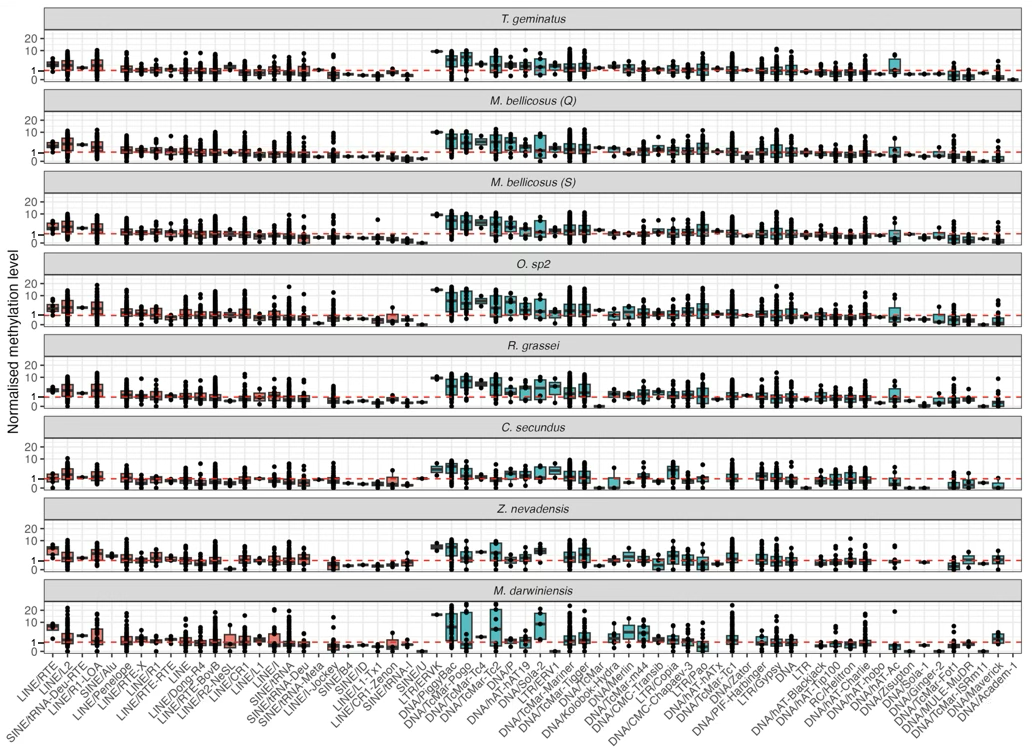

**Supplementary Figure 3.** Boxplots of normalized CpG methylation levels of TE superfamilies for different termite species. TE superfamilies are sorted by TE classes (red for class I TEs and blue for class II TEs) and the mean of the normalized methylation level of the corresponding TE families (points). The normalised methylation level was calculated as the ratio of the percentage of methylated CpG sites in a TE family relative to that of the genome. It ranges from 0 (devoid of methylated CpG sites), 1 (equal to the genomic background; red dashed line), to > 1 (higher than the genomic background). Within each TE superfamily, TE families showed high variation of methylation levels.

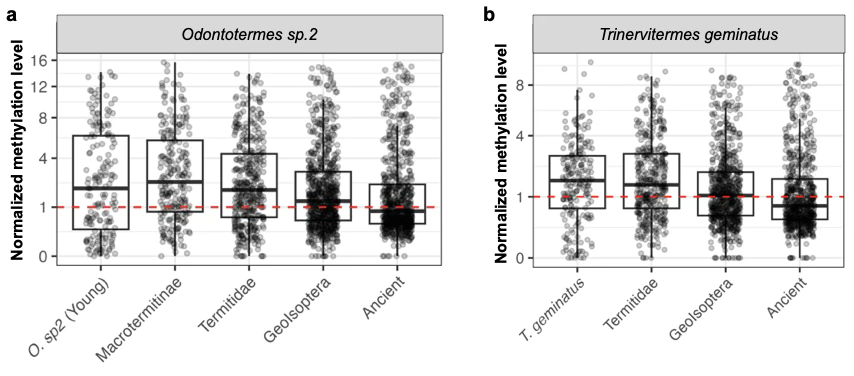

**Supplementary Figure 4.** Boxplots of normalized TE methylation levels across different TE age groups in (**a**) the *Odontotermes* sp.2 and (**b**) the *Trinervitermes geminatus* genome. Red dashed lines represent genomic background.

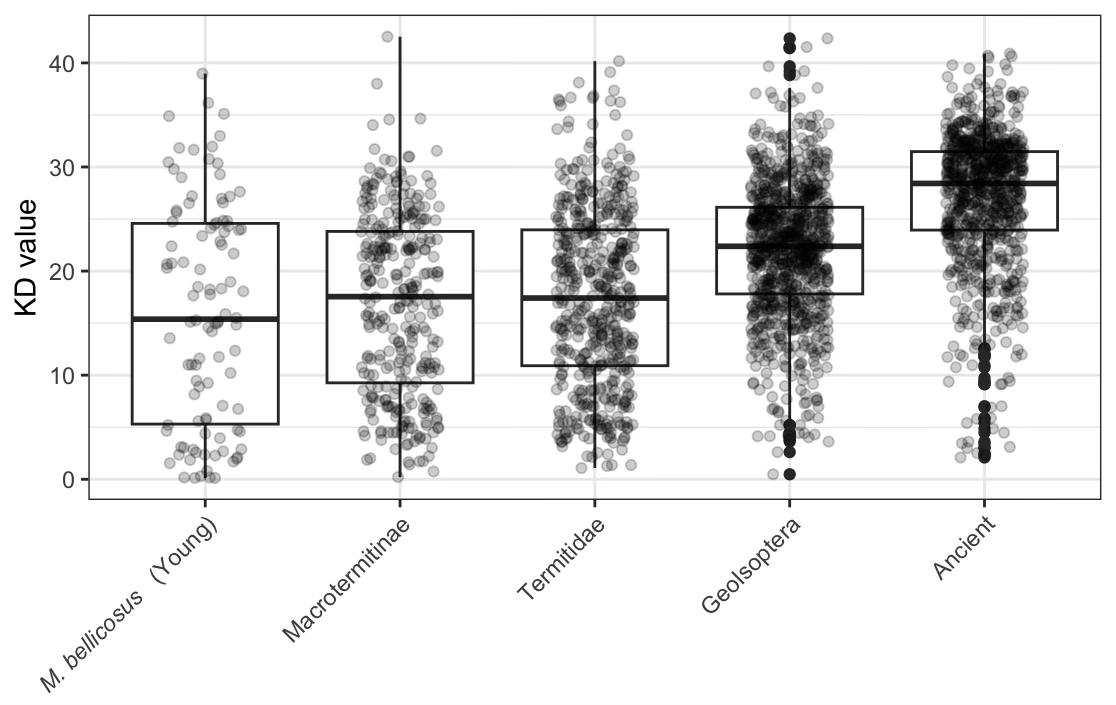

**Supplementary Figure 5. Boxplots of KD values of TE families across different TE age groups**. The mean KD values of ancient TE families were significantly higher than those of younger families (two-sided t-test; *p* < 1e-5), as expected since most ancient TE families are TE remnants.

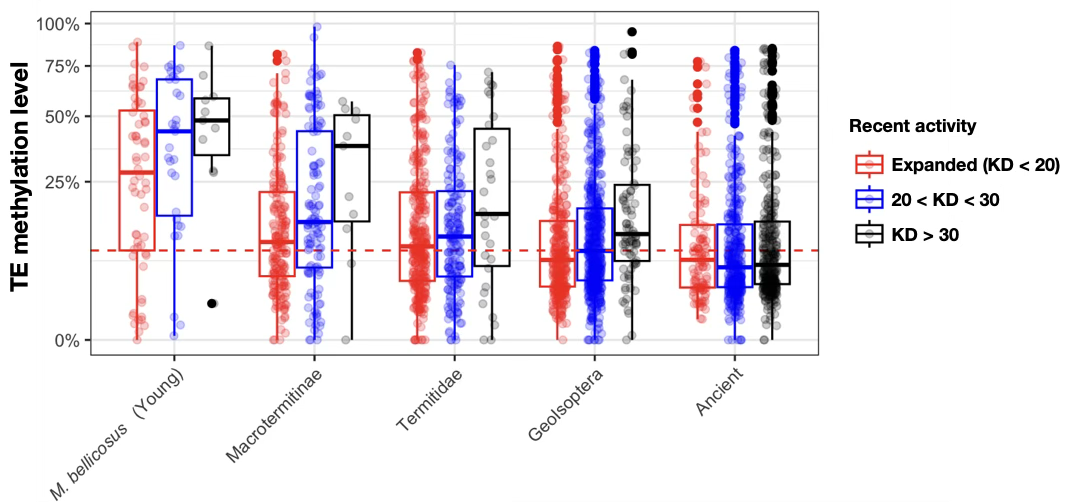

**Supplementary Figure 6. Boxplots of TE methylation levels across different TE age groups**. Within each age group, TE families are stratified into recently expanded (Kimura’s distance, KD < 20; red) and more diverged (indication of non-active) (KD > 20; blue and black) categories. Overall, TE methylation levels declined with increasing TE age. However, more diverged TE families exhibited significantly higher methylation levels than recently expanded families in all age groups, except for the most ancient TEs.

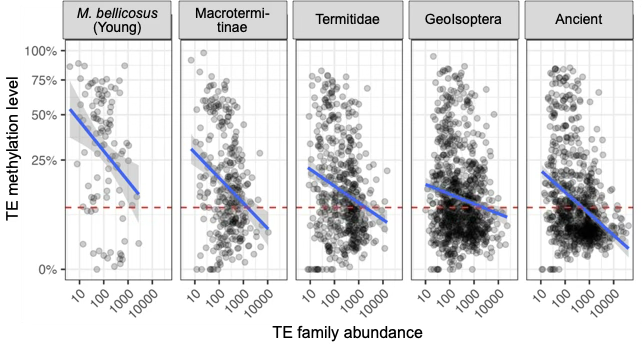

**Supplementary Figure 7.** Association between TE family abundance and TE methylation levels in the *M. bellicosus* genome, plotted separately for each TE age group. Blue lines represent regression lines. The red dashed line represents the genomic methylation level (8%) for *M. bellicosus*.

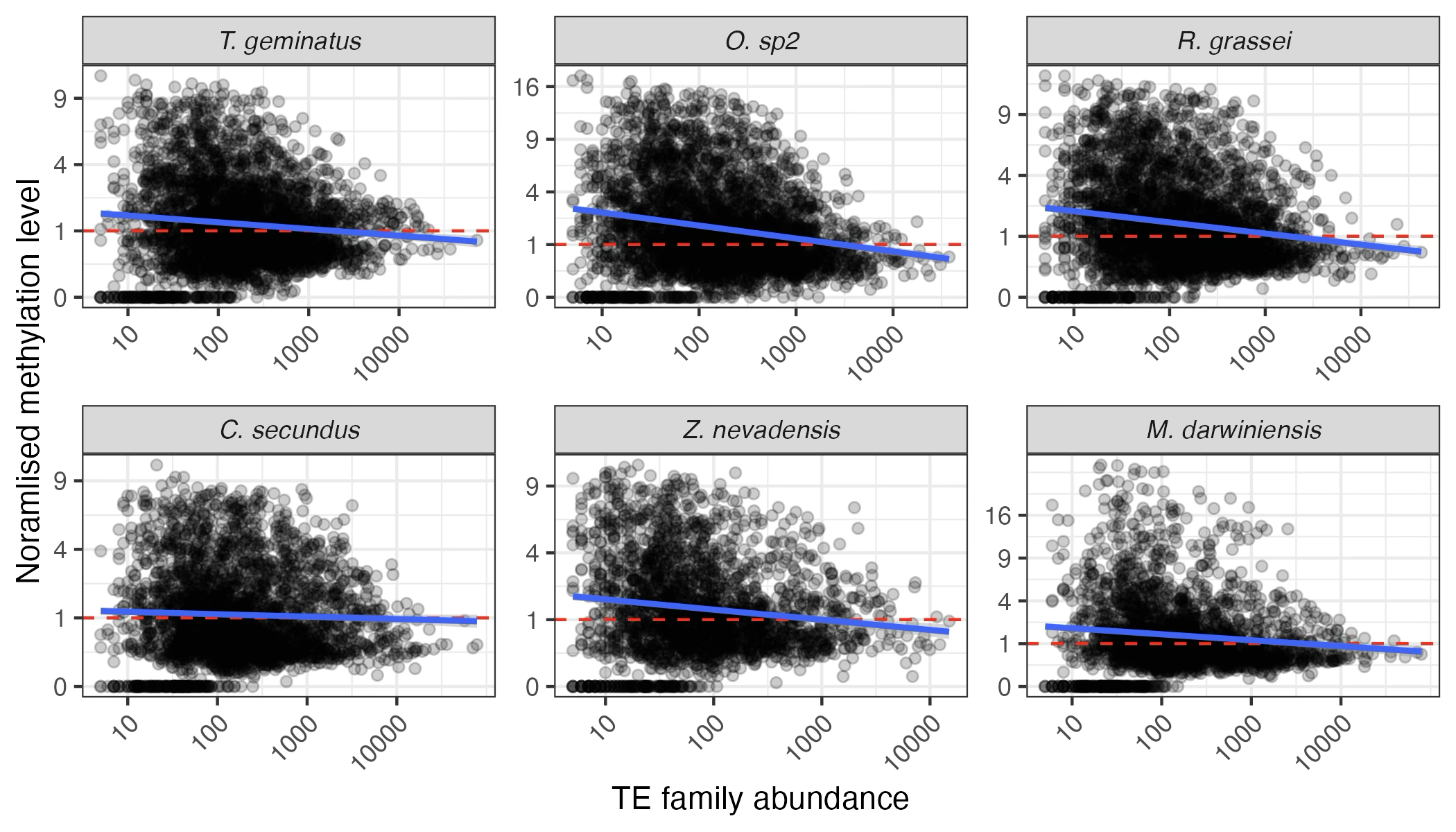

**Supplementary Figure 8.** Association between the normalised methylation levels and TE family abundance, plotted separately for each termite genome. TE methylation and abundance were negatively correlated in all species (Spearman’s correlation test; all *p* < 1e-5, except for *C. secundus*). Blue lines represent regression lines. Red dashed lines represent the genomic backgrounds.

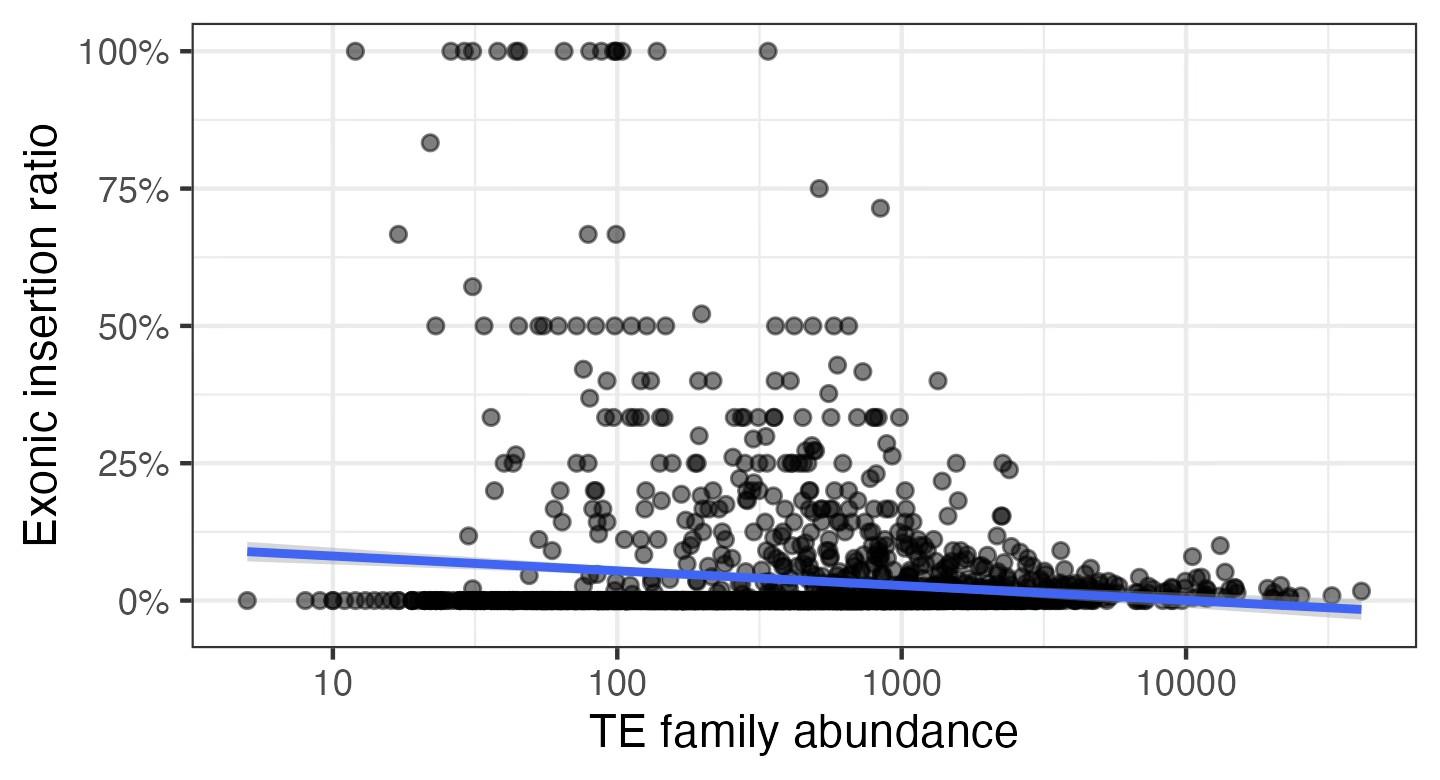

**Supplementary Figure 9.** Association between TE family abundance and TE-associated structural variants (SVs) exonic insertion ratios in *M. bellicosus* genomes. The exonic insertion ratio of a TE family was calculated as the percentage of exonic insertions in relation to the total number of insertions. The blue line represents the regression line. While the majority of TE families had zero exonic insertions, TE families with high abundances were with fewer exonic insertions.

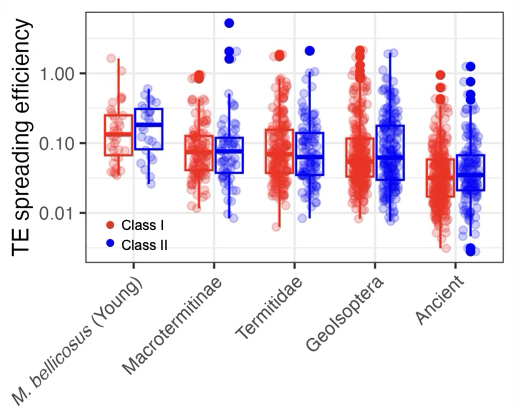

**Supplementary Figure 10.** Boxplots of the TE spreading efficiency across different TE ages in the two TE classes (red for class I TEs and blue for class II TEs). TE spreading efficiency was calculated as the number TE-associated SVs normalised by TE family abundance in the *M. bellicosus* genome.

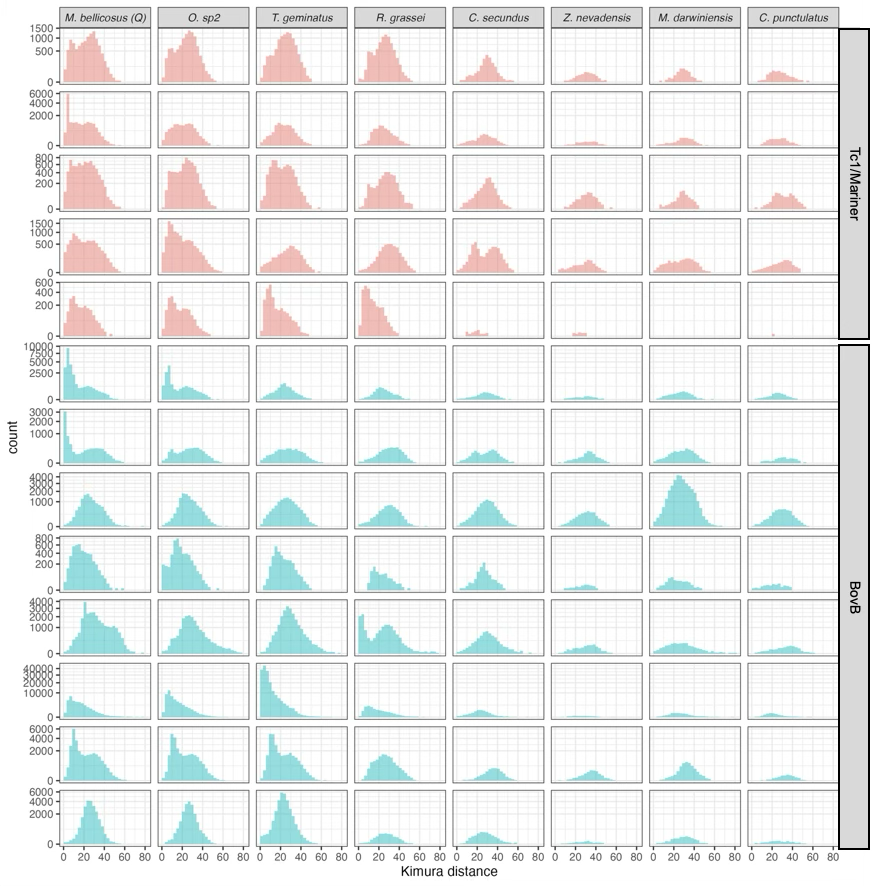

**Supplementary Figure 11.** Histograms of Kimura distances (KD) for the top active Tc1/Mariner (red) and BovB (blue) ancient TE families, plotted separately for each TE family (row) and each species (column). A low KD value indicates recent TE expansion/ invasion, while a high KD value suggests TE remnants that have been inactive for a long time. A combination of both (i.e., a bimodal distribution) suggests that the TE family includes both copies of inactive TE remnants and copies of recently expanded TEs.

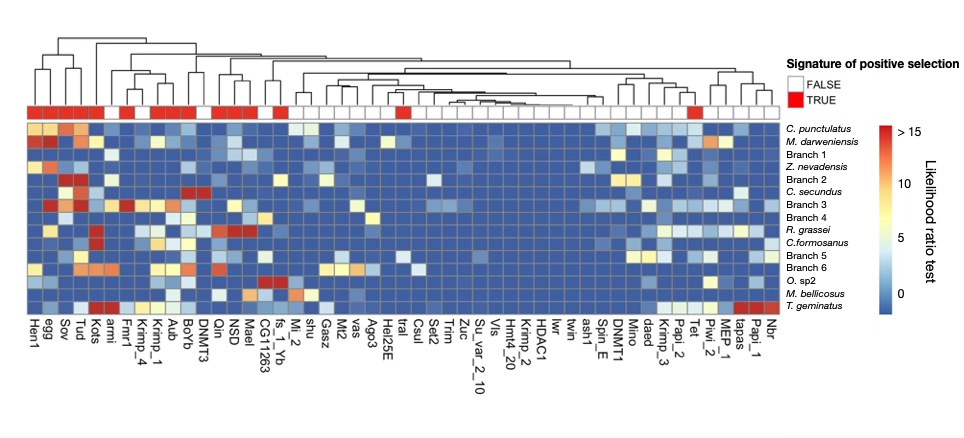

**Supplementary Figure 12**. Heatmap of the signatures of positive selection in TE defense genes in termites and the woodroach. Signatures of positive selection were quantified by likelihood ratio test statistics, ranging from 0 (blue) to > 15 (red). A higher value indicates stronger evidence that the gene has experienced positive selection in the target species or branch. The bar at the top of the heatmap illustrates the overall evidence for positive selection based on all species, with red indicating strong evidence (*p* < 0.05) (multiple testing correction with false discovery rate). To the right of the heatmap are the names of the tested species and branches. Branch 1, 2, 3, 4, 5 and 6 are the branch leading to the last common ancestor (BLCA) of Zootermopsis and the Icoisoptera, the BLCA of Icoisoptera, the BLCA of Geoisoptera, the BLCA of Heterotermitidae, the BLCA of Termitidae, and the BLCA of Macrotermitinae, respectively.

**Supplementary Note 1. Caste effect on TE abundance and TE CpG methylation**

We also looked at the caste difference in TE abundance and TE CpG methylation. Compared to the other termite genomes, the soldier and the queen genome assemblies of *M. bellicosus* were highly similar in their TE abundance and TE methylation at the TE family level (Spearman’s *r* = 0.99 and 0.88, respectively). However, the queen genome had a significantly lower TE abundance and a significantly higher TE methylation level than the soldier genome (two-sided paired Wilcoxon signed-rank test: both *p* < 1e-5). As *M. bellicosus* queens have a high expression level for TE suppressing genes regardless of age ^4^, the TE abundance difference between the queen and the soldier genome might be associated with the lower TE activities during queen ageing.

**Supplementary Note 2. Validation of CpG 5mC identification**

To validate the CpG 5mC identification method, we quantified the CpG methylation levels across different genomic regions, measured by the percentages of methylated CpG sites normalised against the species-specific genomic background. For all species, exonic regions had significantly higher CpG methylation levels than the introns and the genomic background (two-sided binomial test: *p* < 10e-5). Intergenic regions were significantly depleted of methylated CpG sites (two-sided binomial test: *p* < 10e-5) (**Supplementary Fig. A**). Also, CpG methylation levels in 3 prime untranslated regions (3’ UTRs) were significantly higher than in 5’ UTR. These results are consistent with previous findings in *Z. nevadensis* and *B*. *germanica* genomes ^1–3^, demonstrating the validity of using PacBio HiFi reads to identify CpG 5mC in termite genomes.

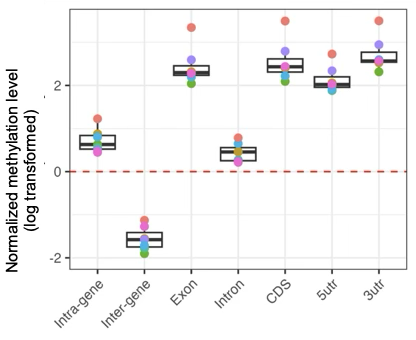

**Supplementary Figure A.** Boxplots of log transformed normalised CpG site methylation levels in different genomic regions in the seven termite genomes. Normalised methylation level was calculated as the ratio of the percentage of methylated CpG sites in the target genomic region to that in the whole genome. After log transformation, a positive value indicates a higher CpG methylation level than the genomic background and a negative value indicates a lower one. Each colour represents one termite species.

**Supplementary Note 3. Associations between TE methylation level and TE abundance at the superfamily level**

TE methylation levels can be positively or negatively associated with TE abundance at the superfamily level across the termite phylogeny (**Supplementary Fig. B**). For example, TE abundance in *Pao*, an LTR retrotransposon superfamily, showed a positive association between methylation level and abundance, whereas *Maverick* and *MULE*, two DNA transposon superfamilies, showed negative associations. Interestingly, while *Pao* comprises a mix of recently expanded (low mean Kimura’s distance; KD < 20) and decayed families (KD > 20), *MULE* and *Maverick* in Geoisoptera consist mostly of recently expanded families that are hypo-methylated (**Supplementary Fig. C**), consistent with the suppressive role of DNA methylation on TE activity. These results suggest that the direction of the association between TE abundance and methylation at the superfamily level may be shaped by the invasion histories of constituent TE families.

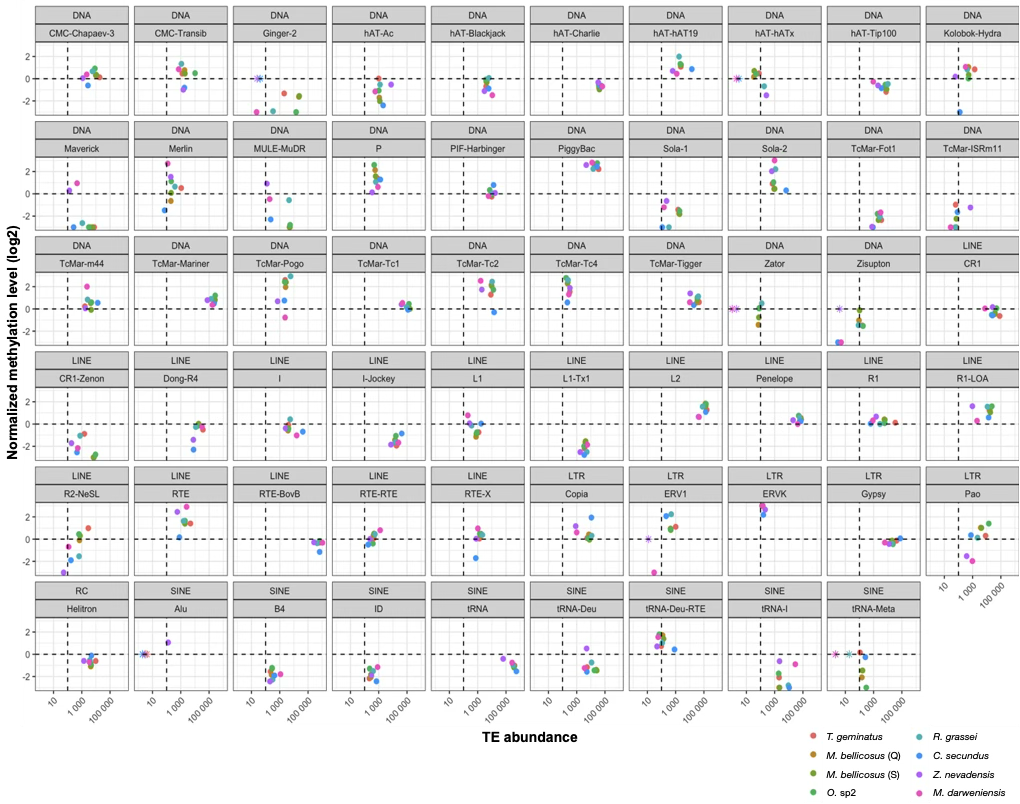

**Supplementary Figure B. Association between TE abundance (x-axis) and normalized TE methylation level (y-axis, log₂ scale) for 59 TE superfamilies across the termite phylogeny.**  For TE superfamilies that were species-specific (e.g., Alu elements in *Z. nevadensis*), normalized DNA methylation values were not available in other species and were thus set to zero; these TE superfamilies are marked with an asterisk (*). The horizontal dashed line represents the normalized genomic methylation level (0) and the vertical dashed line represents 100 TE copies.

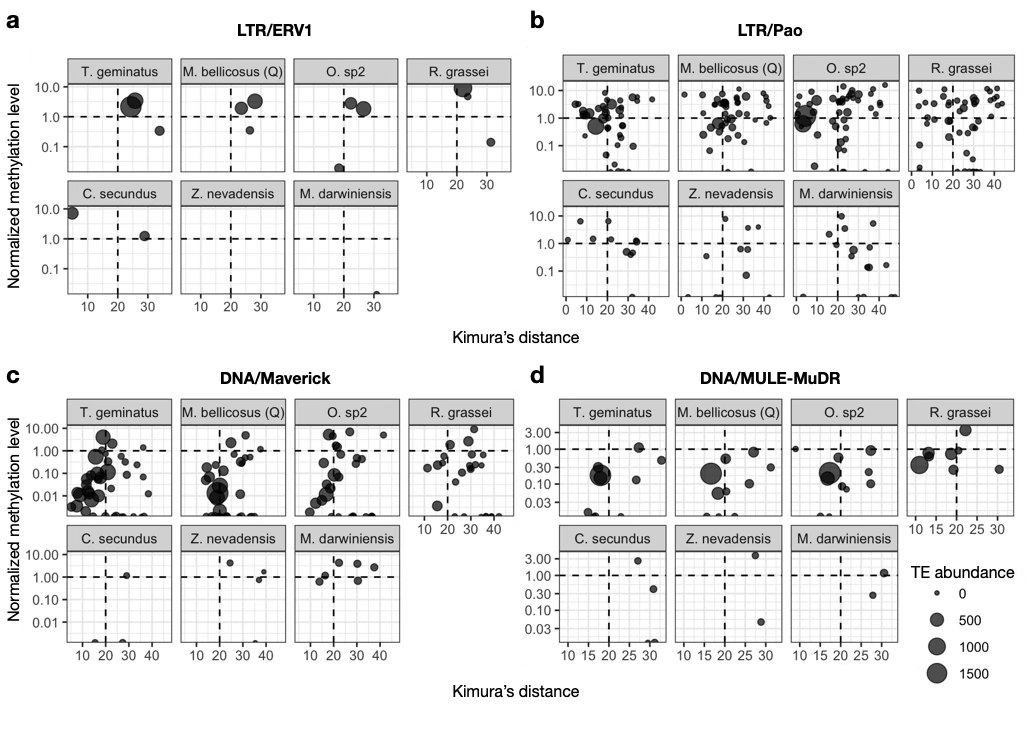

**Supplementary Figure C.** ERV1 and Pao (panel **a** and **b**), two LTR retrotransposon superfamilies with high DNA methylation levels in species with high TE abundance, consist either of decayed TE families (Kimura’s distance, KD > 20) or a mixture of recently expanded (KD < 20) and decayed TE families. In contrast, Maverick and MULE (panel **c** and **d**), two DNA transposon superfamilies with low DNA methylation levels in species with high TE abundance, consist primarily of recently expanded TE families, consistent with the suppressive role of DNA methylation on TE activity. These patterns suggest that the relationship between TE abundance and methylation at the superfamily level may reflect the evolutionary histories of the underlying TE families. Each dot represents one TE family, with the dot size, x value, and y value indicating the abundance, KD, and normalized methylation level of the TE family, respectively.

**Supplementary Note 4. Inferring active TE families from TE derived structure variants (SVs)**

TE-derived SVs can arise from both active TEs and inactive TEs that have decayed and been lost in some populations. By quantifying the distribution of SV insertions (relative to the reference genome) across five individual termite samples (10 haplotypes), we found that most insertions occurred in only one or two haplotypes (**Supplementary Fig. D**). Because loss of a TE in the reference genome would result in the same “insertion” being detected across multiple haplotypes, the predominance of haplotype-specific insertions suggests that these SVs represent true TE insertions rather than decayed TE sequences missing from the reference genome.

In contrast, for SV deletions it is more difficult to distinguish between true TE excisions (cut-and-paste events) and TE loss, since both can generate haplotype-specific deletions.

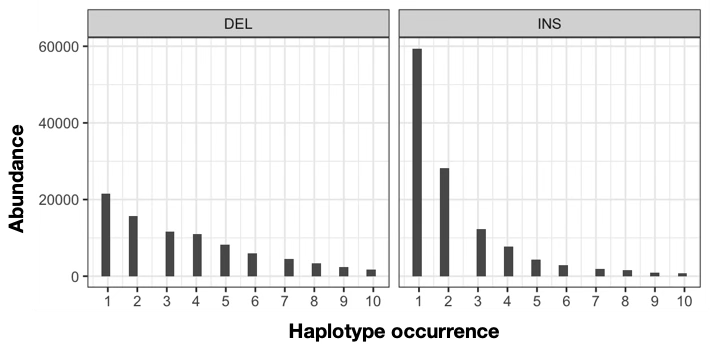

**Supplementary Figure D. Haplotype frequency distribution of SV deletion (left) and SV insertion (right) across the five individual termite samples comparing to the reference genome.** The majority of SV insertions occur only in one or two haplotypes, indicating these SVs are derived from active TEs.

**Supplementary Note 5. Contribution of TE superfamilies to genome size variation among termites**

We sorted TE superfamilies by their size variation among termite genomes and found that RTE-BovB (LINE), tRNA (SINE), Gypsy (LTR retrotransposon), TcMar-Mariner (DNA transposon), L2 (LINE), and Tc1/mariner (DNA transposon) show the highest variation. Notably, BovB and Tc1/mariner are also among the top three active TE superfamilies in *M. bellicosus* (together with Ginger-2, a DNA transposon that also exhibits high size variation). This suggests that genome size variation among termites can be partly attributed to TE families that remain active.
